# High culturable diversity and climate-associated seasonal dynamics of Saccharomycotina yeasts in subtropical forest leaf litter

**DOI:** 10.64898/2026.08.25.747014

**Authors:** Wei-Ting Chien, Yu-Chen Yeh, Cheng-Ju Yang, Yu-Ching Liu, Chen Hsiao, Pei-Wei Sun, Chia-Hsiu Tsai, Po-Ju Ke, Chau-Ti Ting, Chia-Hao Chang-Yang, Isheng Jason Tsai

## Abstract

Forest-associated Saccharomycotina occur at low relative abundance, limiting inference about their diversity and dynamics. We sampled leaf litter weekly for 47 weeks across a subtropical forest in northern Taiwan. Enrichment, isolation and ITS sequencing recovered 687 isolates, including 613 Saccharomycotina representing 56 described species and 77 putatively novel operational taxonomic units. Rarefaction indicated unsampled culturable diversity. Among litter traps, community dissimilarity was high and dominated by taxon replacement, but neither topography nor geographic distance was associated with composition, and turnover matched randomised expectations. Richness peaked during warm, wet periods and declined in winter, and minimum temperature showed the strongest statistical association. Composition was associated with maximum temperature, minimum relative humidity, precipitation and solar radiation. Selected isolates’ thermal optima covaried with collection-week temperatures, and two October *Magnusiomyces magnusii* isolates had higher optima than four winter isolates. Together, these findings reveal substantial culturable diversity and seasonal community restructuring consistent with temperature-related filtering.

## Introduction

Saccharomycotina yeasts are widely studied in biotechnology, industry and fundamental research. The brewer’s yeast *Saccharomyces cerevisiae* underpins baking and brewing, has applications in biofuel production, pharmaceuticals and synthetic biology, and is a major model for eukaryotic cell biology^1^. Other Saccharomycotina species also have biotechnological potential^2–4^. Despite this importance, their diversity and ecological roles in natural environments remain poorly understood^5,6^, partly because yeasts generally occur at low relative abundance^7^. In temperate forest ecosystems, yeast sequences comprise only 0.4–14.3% of fungal reads in soil and 0.2–9.9% in leaf litter, and these sequences are dominated by Basidiomycota, with Saccharomycotina comparatively scarce^8^. *Saccharomyces cerevisiae,* for example, has an estimated average relative abundance of only 0.012% in environmental samples^9^. This scarcity limits estimates of natural diversity from environmental sequencing and has reinforced the historical focus on economically or clinically important species^10–12^.

Beyond incomplete inventories, how Saccharomycotina communities are structured across local space and through time remains unclear^7^. Broad-scale analyses associate yeast distributions with vegetation, topography and precipitation^5,13^, while whole-fungal surveys of Fagaceae forests have documented substantial turnover among neighbouring habitats and between seasons^14^. Field distributions of wild *Saccharomyces* and the abundance of oak-associated fermentative yeasts are associated with climatic conditions, particularly temperature, while thermal responses differ among species, populations from contrasting climates and geographically separated isolates^15–17^. However, these relationships have largely been examined across taxa or broad geographic gradients. Combining spatially replicated time-series sampling with targeted thermal phenotyping can therefore distinguish persistent spatial heterogeneity from temporal community reorganisation and test whether seasonal occurrence covaries with thermal physiology.

Fagaceae forests provide a tractable system for addressing these questions because wild Saccharomycotina are repeatedly recovered from oaks and associated forest substrates^6,18^. A recent survey across 28 *Quercus* forests found communities dominated by *Saccharomyces*, *Kluyveromyces* and *Pichia*, with marked geographical variation and associations with temperature and precipitation^16^. Other genera associated with *Quercus* include *Lachancea* and *Hanseniaspora*^19^. Taiwan provides an opportunity to extend these observations to community-level ecology. Forests cover 60.7% of the island^20^, broadleaf forests account for 67% of forest area, and natural vegetation forms pronounced altitudinal belts that include *Castanopsis*- and *Quercus*-rich forests. Extensive surveys of Taiwanese broadleaf forests have also recovered nine lineages of *S. cerevisiae*, including three endemic lineages, with genetic diversity comparable to that sampled across continental Asia and distinct lineages coexisting on individual trees^9,21^. Taiwanese Fagaceae forests therefore combine documented wild-yeast diversity with pronounced environmental heterogeneity, providing a suitable system for resolving community structure across space and time.

To test whether spatial heterogeneity and climatic variation were associated with culturable Saccharomycotina communities, we sampled leaf litter over 47 weeks across two warm-to-winter intervals at the Fushan Forest Dynamics Plot (FDP), a Fagaceae-dominated evergreen forest in northern Taiwan. Weekly samples from 20 spatially distributed traps were processed by selective enrichment and ITS sequencing. We tested whether geographic distance, altitude, slope and distance to the nearest stream accounted for among-trap composition^8,14^. We also tested whether yeast richness and composition varied seasonally with temperature- and moisture-related conditions^19,22^. Targeted thermal phenotyping assessed whether field recovery temperature corresponded to variation in thermal physiology.

## Results

### Sampling effort and yeast recovery

We collected leaf litter weekly from 20 traps across the 500 × 500 m Fushan Forest Dynamics Plot in northern Taiwan (**Fig. 1**). Sampling spanned October 2022 to mid-February 2023 (19 weeks) and August 2023 to mid-February 2024 (28 weeks), yielding 940 litter samples and 1,880 parallel incubations at 20°C and 12°C. Yeasts were recovered from 486 samples, with growth detected in 643 incubations. These cultures yielded 687 isolates, including 264 from the first period and 423 from the second. Colony PCR and ITS sequencing identified 232 of 264 isolates (87.9%) from the first period. During the second period, ITS3/ITS4 was used when full-length ITS amplification failed, increasing identification to 94.6% (400 of 423 isolates).

**Fig. 1.**
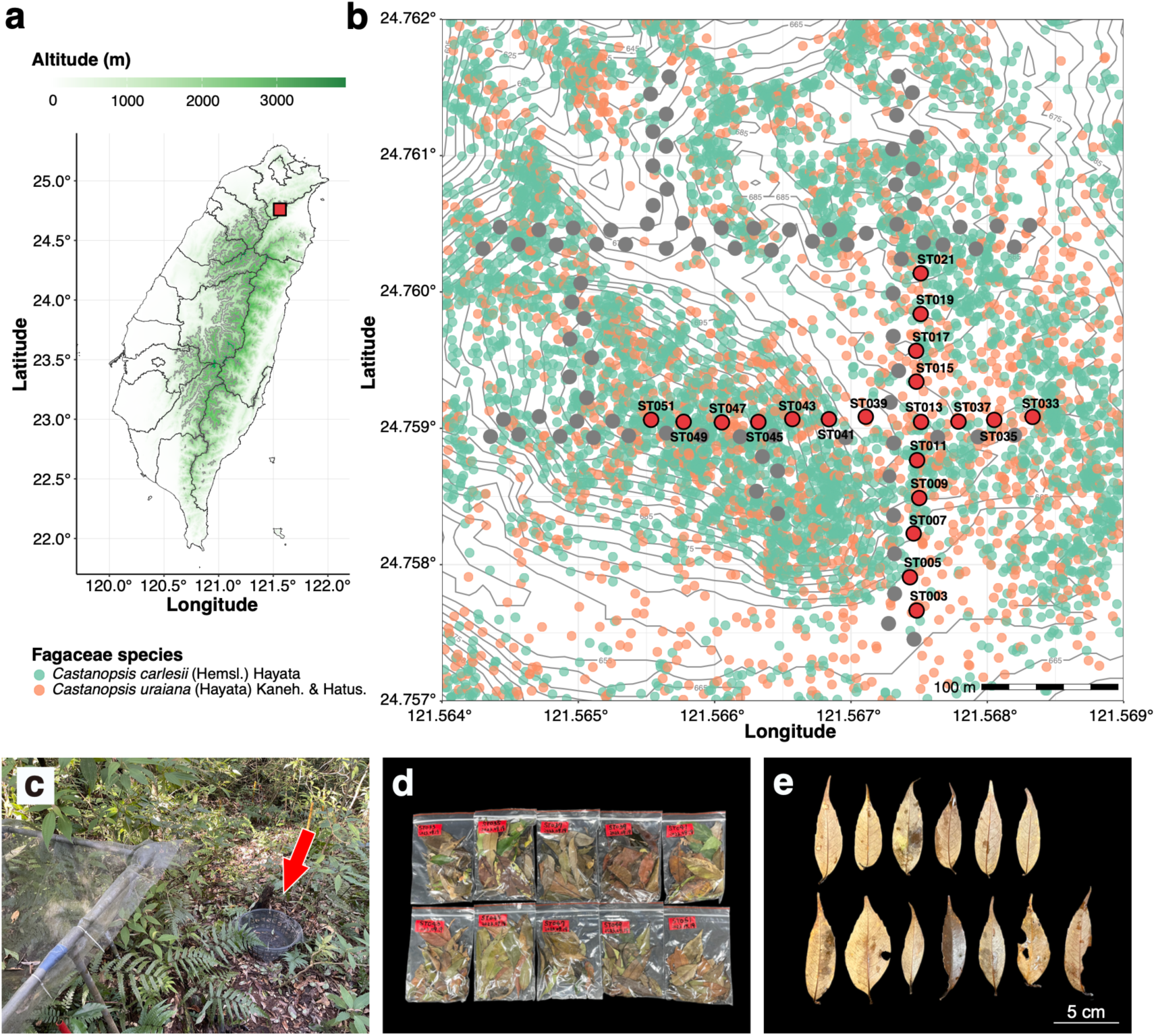
Study site, sampling layout and leaf-litter material. (a) Location of the Fushan Forest Dynamics Plot (FDP) in northern Taiwan. The red square marks the plot and shading denotes altitude. (b) Topographic map of the 500 × 500 m plot. Grey and red circles mark permanent seed traps and the 20 litter traps sampled here, respectively. Green and orange points mark *Castanopsis carlesii* and *Castanopsis uraiana*, respectively. (c) Litter trap used for weekly collection. (d) Representative weekly samples. (e) Representative Fagaceae leaf pieces used for enrichment.

Across both periods, 632 isolates were identified (92.0%). Of these, 613 belonged to Saccharomycotina, comprising 56 described species and 77 putatively novel operational taxonomic units (OTUs; **Supplementary Tables S1 and S2**). To minimise primer-dependent bias, ecological analyses included only ITS1Fngs/ITS4 identifications. Repeated recovery of a taxon within a trap × week sample was scored once, yielding 497 trap × week–taxon incidence records across 124 taxa, comprising 56 described species and 68 putatively novel OTUs (**Supplementary Table S2**).

Based on 497 incidence records across 124 taxa in the primer-standardised ecological dataset, the asymptotic richness estimate was approximately 351 taxa, suggesting that substantial culturable diversity remained undetected (**Fig. 2a**). Of the 124 taxa, 77 (62.1%) were detected in only one trap × week sample. Putatively novel OTUs had fewer incidence records per taxon than described species (Wilcoxon rank sum test, P < 0.001; **Fig. 2b**, inset). The Saccharomycotina assemblage was therefore characterised by many rarely detected taxa.

**Fig. 2.**
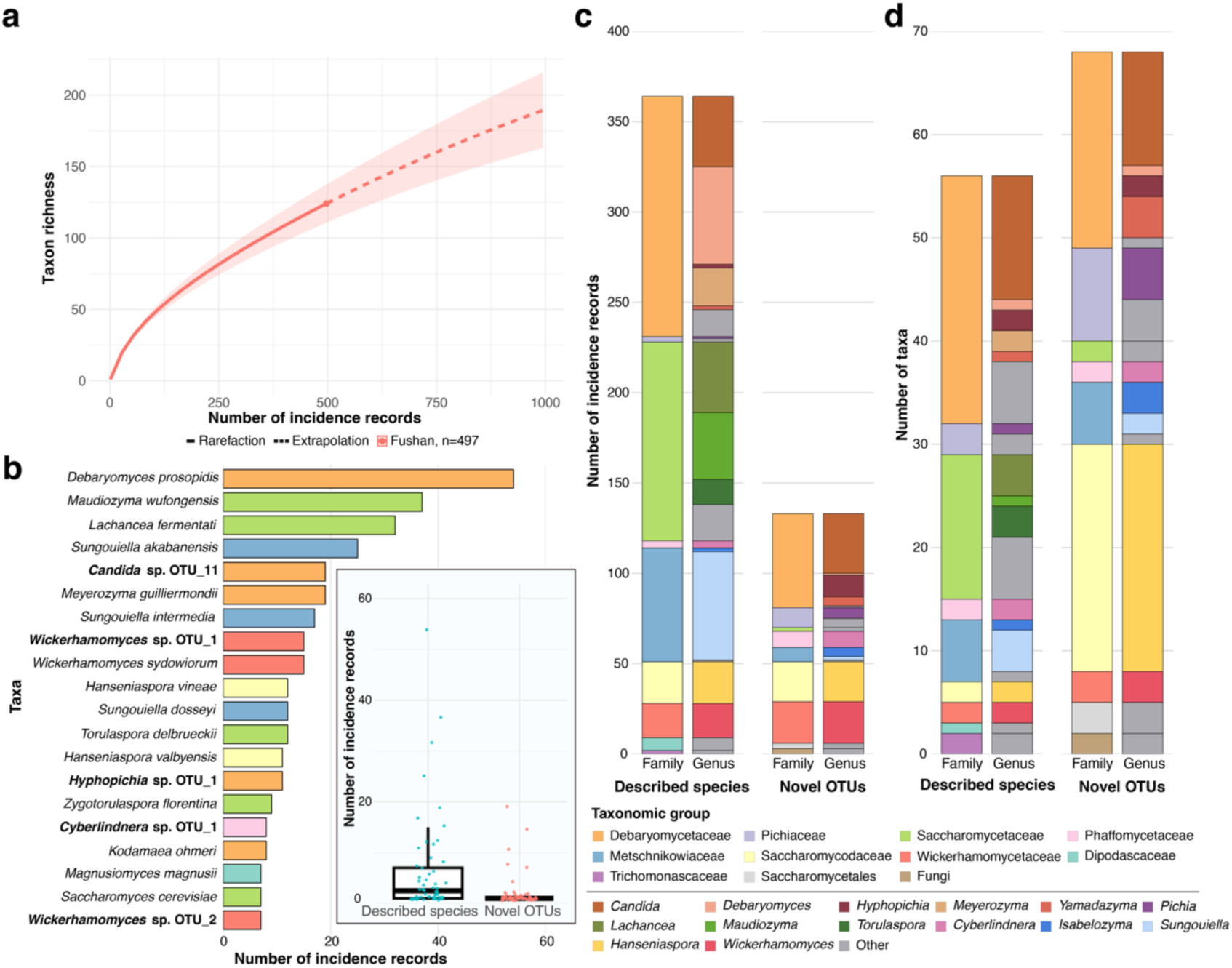
Yeast recovery and taxonomic composition of the primer-standardised ecological dataset. (a) Record-based rarefaction (solid line) and extrapolation (dashed line) of taxon richness. The point marks the observed endpoint of 497 incidence records across 124 taxa, and shading denotes the 95% confidence interval. (b) The 20 taxa with the most incidence records. Bold labels denote putatively novel OTUs, and the inset compares per-taxon incidence records between described species and putatively novel OTUs. (c) Incidence records and (d) taxon richness across families and genera, separated into described species and putatively novel OTUs. Colours identify taxonomic groups.

### Taxonomic composition of dominant and novel lineages

Within the primer-standardised ecological dataset, the most frequently detected described species was *Debaryomyces prosopidis* (54 incidence records), followed by *Maudiozyma wufongensis* (37 records) and *Lachancea fermentati* (32 records), each detected in more than 30 trap × week samples (**Fig. 2b**). Several fermentation-associated yeasts were also detected repeatedly, including *Hanseniaspora valbyensis* (11 records), *H. vineae* (12 records), *Saccharomyces cerevisiae* (seven records) and *Torulaspora delbrueckii* (12 records; **Fig. 2b**). The opportunistic pathogens *Kodamaea ohmeri* and *Candida albicans* were detected in eight (**Fig. 2b**) and four (**Supplementary Table S5**) trap × week samples, respectively. Among novel OTUs, *Candida* sp. OTU_11 was the most frequently detected lineage (19 records), followed by *Wickerhamomyces* sp. OTU_1 (15 records) and *Hyphopichia* sp. OTU_1 (11 records).

At the family level, Debaryomycetaceae was the most frequently detected family, comprising 185 incidence records, including 133 from described species and 52 from novel OTUs (**Fig. 2c**). The family included multiple genera, including *Candida*, *Debaryomyces*, *Hyphopichia*, *Meyerozyma* and *Yamadazyma*. *Candida* accounted for 72 records across 12 described species (39 records) and 11 novel OTUs (33 records), representing 23 taxa in total. *Debaryomyces* contributed 55 records, of which 54 were attributed to *D. prosopidis*. By contrast, Saccharomycetaceae comprised 112 records, including 110 from 14 described species and only two from two novel OTUs. This family included the genera *Lachancea*, *Maudiozyma*, *Torulaspora* and *Saccharomyces*. Pichiaceae, although represented by only 14 incidence records, was dominated by novel OTUs, with 11 records from nine novel OTUs and three records from three described species. At the genus level, *Hanseniaspora* showed the greatest taxonomic richness, comprising two described species and 22 novel OTUs (**Fig. 2d**). The 22 novel OTUs were each represented by a single incidence record, whereas the two described species together accounted for 23 records.

To determine whether the 22 novel *Hanseniaspora* OTUs were concentrated within one lineage or distributed across the genus, we constructed an ITS phylogeny. Sixteen OTUs formed a well-supported clade near *H. vineae*, three were placed near *H. valbyensis* and three clustered with *H. meyeri* and *H. uvarum* (**Supplementary Fig. 1**). Thus, novel ITS diversity spanned three *Hanseniaspora* lineages.

### Yeast communities show high turnover without detectable spatial structuring

Weekly incidence records were pooled for each of the 20 litter traps across the 47-week survey to characterise among-trap variation in yeast diversity. Each trap was sampled for 47 weeks, and the number of weeks with at least one Saccharomycotina detection ranged from 12 to 29 among traps. Observed richness ranged from 10 to 23 taxa per trap (mean = 15.3), while incidence-based Chao2 estimates ranged from 16.8 to 181.6 taxa (median = 46.1, mean = 54.8; **Fig. 3a**). The highest Chao2 estimate exceeded the 124 taxa detected across the complete trap dataset because of its sensitivity to taxa detected in only one or two sampling weeks and should not be interpreted as observed local richness. Shannon diversity calculated from weekly detection frequencies ranged from 1.94 to 2.94 (median = 2.50, mean = 2.50; **Fig. 3a**). Community composition was analysed using a dbRDA of binary Jaccard dissimilarity calculated from the 20 × 124 trap-by-taxon presence–absence matrix and constrained by altitude, slope and distance to the nearest stream (**Supplementary Table S3**). The model accounted for 16.5% of the total variation but only 0.9% after adjustment (R² = 0.165, adjusted R² = 0.009) and was not significant (pseudo-*F* = 1.06, *P* = 0.30; **Fig. 3b**). Neither altitude (marginal R² = 0.05, *P* = 0.66), slope (marginal R² = 0.05, *P* = 0.44) nor distance to the nearest stream (marginal R² = 0.05, *P* = 0.55) accounted for significant variation. The first two constrained axes represented 6.8% and 5.4% of the total variation, respectively. Across the sampling extent, geographic distance was not associated with Jaccard dissimilarity across all taxa (Mantel r = 0.04, P = 0.32) or recurrent taxa (Mantel r = 0.11, P = 0.18), providing no evidence of monotonic distance decay.

**Fig. 3.**
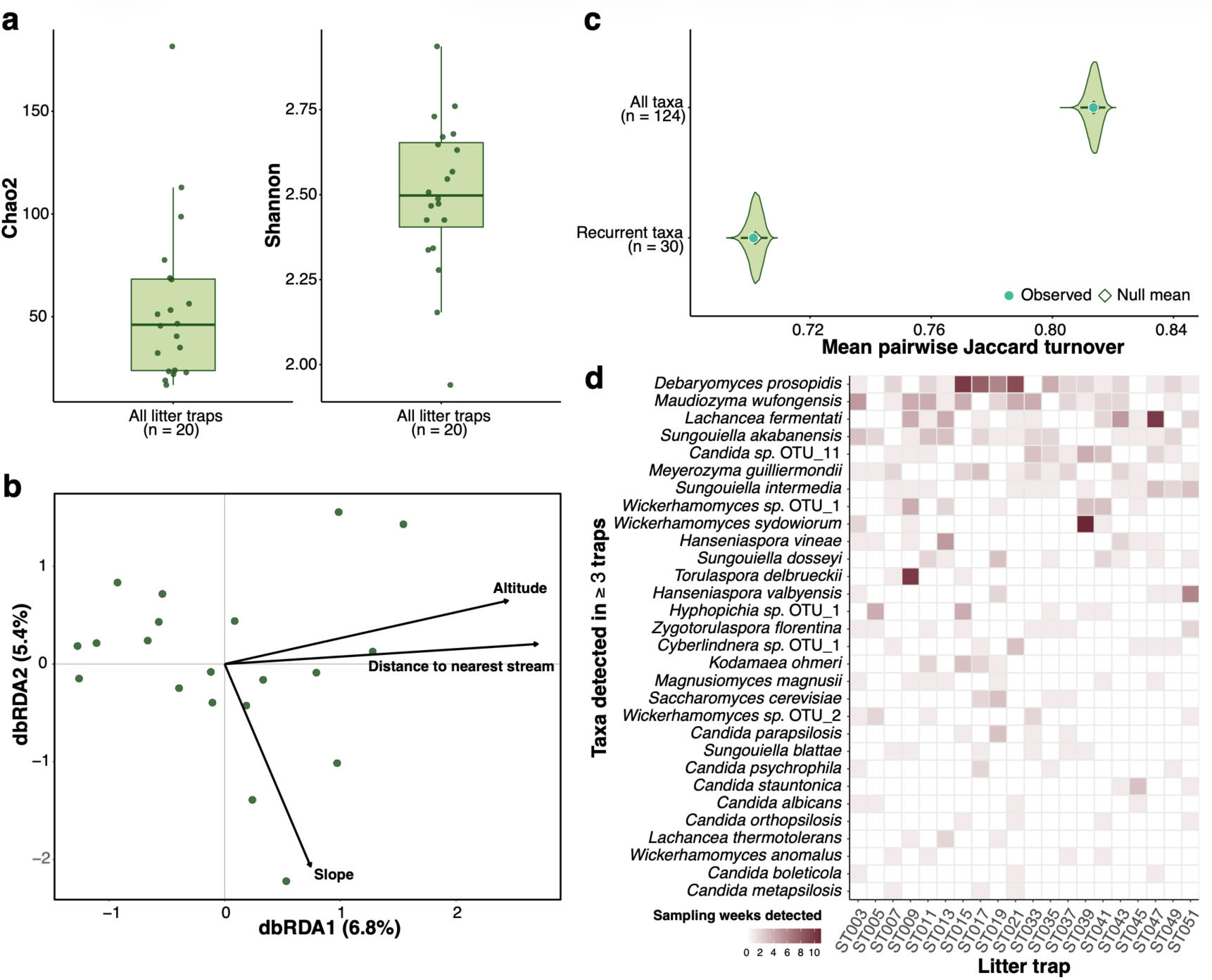
Spatial diversity and community dissimilarity among litter traps. (a) Incidence-based Chao2 richness and Shannon diversity calculated from weekly detection frequencies for the 20 traps. Points denote traps, boxes the interquartile range, centre lines the median and whiskers 1.5 times the interquartile range. (b) dbRDA of binary Jaccard dissimilarity for the trap-by-taxon matrix constrained by altitude, slope and distance to the nearest stream. Points denote traps, arrows fitted predictors and axis labels the proportion of total variation represented. The overall model and marginal terms were assessed with 9,999 unrestricted permutations. (c) Observed mean pairwise Jaccard turnover and null distributions from 10,000 matrices preserving trap richness and taxon occurrence, shown for all 124 taxa and the 30 taxa detected at three or more traps. Filled circles denote observed values and diamonds null means. (d) Occurrence-frequency heatmap for the 30 recurrent taxa. Cell colour gives the number of weeks in which each taxon was detected at each trap. Traps are ordered numerically and taxa by total incidence records.

Despite the absence of detected topographic associations or monotonic distance decay, among-trap dissimilarity remained high. Mean pairwise Jaccard dissimilarity across all 124 taxa was 83.9% (range = 66.7%–96.9%). Turnover and nestedness contributed 81.4 and 2.6 percentage points, respectively, with turnover accounting for approximately 97% of total dissimilarity. Randomisation of the complete incidence matrix showed that total dissimilarity did not differ from its null expectation (observed = 83.9%; null mean = 84.1%, 95% interval = 83.8%–84.3%; two-sided P = 0.304). Turnover was also consistent with its null distribution (observed = 81.4%, null mean = 81.4%, 95% interval = 80.9%–81.8%, two-sided *P* = 0.928) (**Fig. 3c**). Nestedness likewise did not differ from its null expectation (observed = 2.6%, null mean = 2.7%, 95% interval = 2.5%–3.0%, two-sided *P* = 0.336). Within the recurrent-taxon subset, total dissimilarity did not differ from its null expectation (observed = 73.7%, null mean = 73.8%, 95% interval = 73.5%–74.0%, two-sided *P* = 0.790). Turnover was also consistent with its null distribution (observed = 70.1%, null mean = 70.2%, 95% interval = 69.7%–70.6%, two-sided *P* = 0.751) (**Fig. 3c**). Nestedness likewise did not differ from its null expectation (observed = 3.6%, null mean = 3.6%, 95% interval = 3.3%–3.9%, two-sided *P* = 0.787). Turnover nevertheless accounted for 95.1% of recurrent-taxon dissimilarity and remained the dominant component when recurrent taxa were defined as occurring at two, three or five traps (96.0%, 95.1% and 91.7%, respectively). The incidence-frequency heatmap showed the occurrence patterns underlying this high dissimilarity (**Fig. 3d**). Of the 124 taxa, 82 (66.1%) were detected at only one trap, 12 (9.7%) at two traps and 30 (24.2%) at three or more traps. Among-trap dissimilarity was therefore dominated by turnover, but taxon replacement did not differ from randomised expectations after accounting for trap richness and taxon occurrence.

### Temporal variation in yeast diversity

In the primer-standardised ecological dataset, weekly incidence records from the 20 traps were pooled by sampling week across the 47-week survey. Yeasts were recovered throughout both periods, with 1–18 Saccharomycotina-positive samples per week (mean = 8.51). Recovery was highest on 6 September 2023, when 18 of 20 samples were positive, but only one sample was positive on each of 26 December 2023 and 2 January 2024. Observed richness averaged 8.28 and ranged from 1 to 19 taxa per week. Weekly Chao2 estimates averaged 27 (median = 25.1) and ranged from 1 to 91.1, with the maximum recorded on 11 October 2022 (**Fig. 4a**). Shannon diversity showed a similar pattern, declining to zero on 26 December 2023 and 2 January 2024 and reaching 2.84 on 6 September 2023 (median = 2.04, mean = 1.87; **Fig. 4b**).

**Fig. 4.**
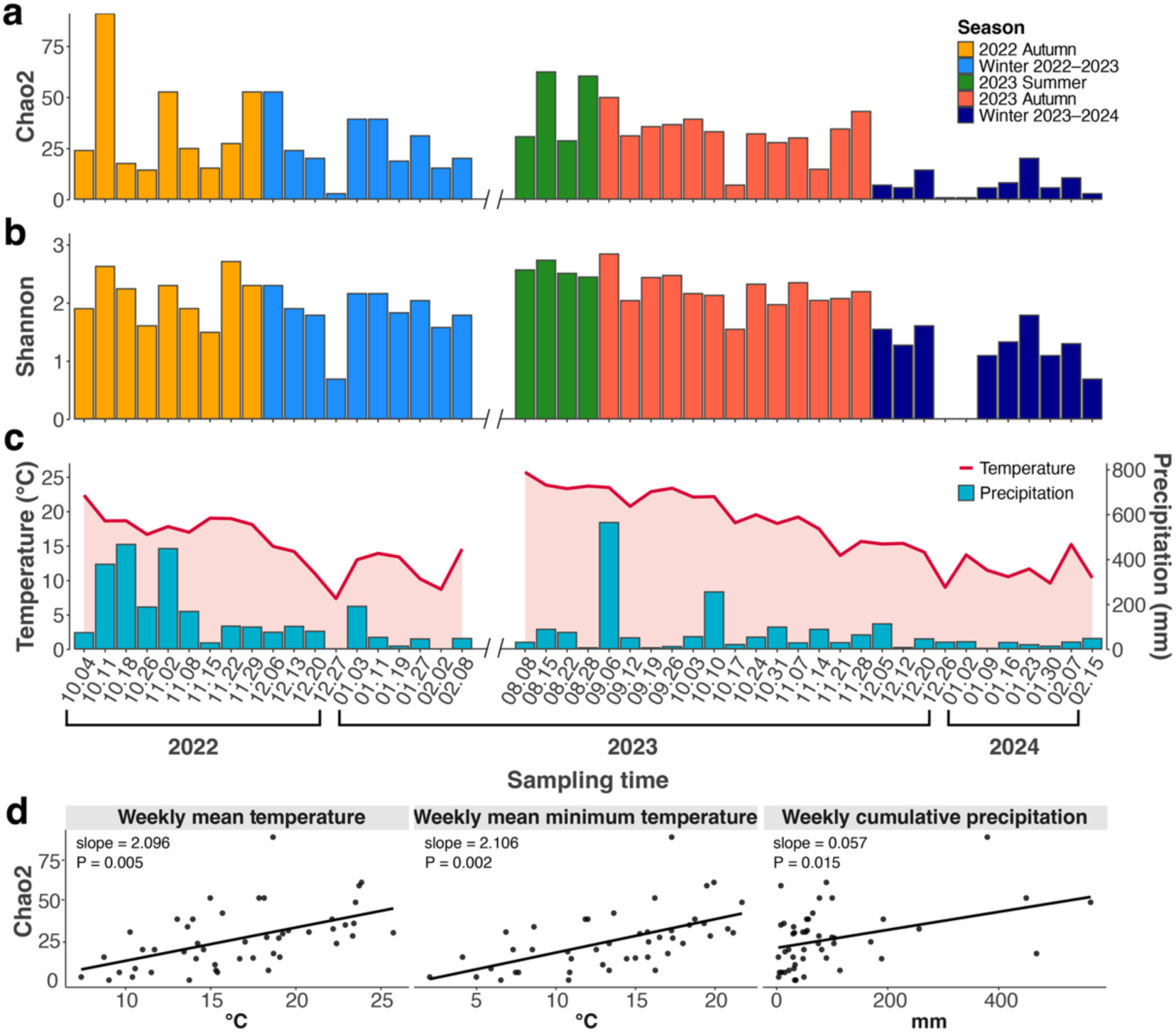
Weekly yeast diversity and climatic conditions. (a) Incidence-based Chao2 richness and (b) Shannon diversity across 47 sampling weeks. Colours distinguish meteorological seasons within each sampling period. (c) Weekly mean temperature (line) and cumulative precipitation (bars). Breaks in (a–c) separate the sampling periods. (d) Relationships between Chao2 richness and weekly mean temperature, mean minimum temperature and cumulative precipitation. Points represent sampling weeks, and lines show fixed-effect fits from separate linear mixed-effects models with year–month as a random intercept. Panel annotations give estimated slopes and P values. Relationships with the complete set of climatic variables are shown in **Supplementary Fig. 2**.

Diversity was generally higher during summer and autumn and lower during winter (**Figs. 4a–c**). Of the climatic variables tested, weekly mean minimum temperature showed the strongest association with Chao2 richness (*P* = 0.002). Richness was also positively associated with weekly mean temperature (*P* = 0.005), mean maximum temperature (*P* = 0.015) and cumulative precipitation (*P* = 0.015; **Fig. 4d**; **Supplementary Fig. 2**; **Supplementary Table S3**). No significant associations were detected with mean or minimum relative humidity, mean wind speed, maximum gust speed or global solar radiation.

### Temporal shifts in community composition

A dbRDA of binary Jaccard dissimilarity among weeks tested whether climatic conditions were associated with temporal community composition (**Supplementary Table S3**). The model accounted for 15.8% of total variation (R² = 0.158) but 6.4% after adjustment (adjusted R² = 0.064). The overall model was significant (pseudo-F = 1.688, P = 0.001; **Fig. 5a**). Marginal permutation tests detected significant associations with maximum temperature (marginal R² = 0.042, P = 0.001), minimum relative humidity (marginal R² = 0.033, P = 0.038), cumulative precipitation (marginal R² = 0.034, P = 0.020) and global solar radiation (marginal R² = 0.039, P = 0.005). The first two constrained axes represented 5.7% and 3.9% of total variation, respectively. Along dbRDA1, winter communities occurred primarily on the negative side and aligned with higher minimum relative humidity, whereas summer and autumn communities occurred mainly on the positive side and aligned with higher maximum temperature, global solar radiation and precipitation.

**Fig. 5.**
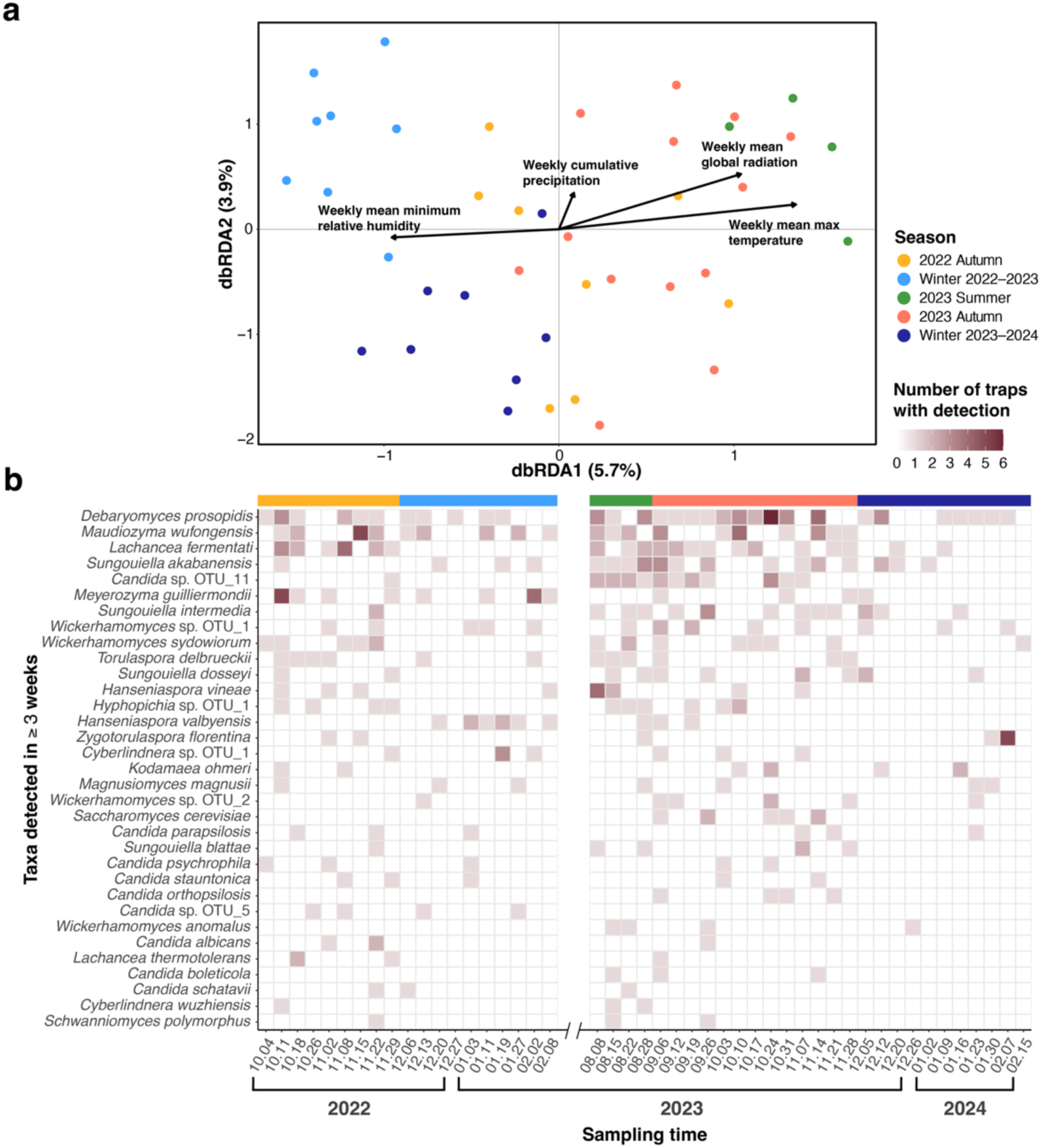
Temporal differentiation of yeast communities and recurrent taxa. (a) dbRDA of binary Jaccard dissimilarity among weekly communities constrained by maximum temperature, minimum relative humidity, cumulative precipitation and global solar radiation. Points denote weeks and colours meteorological seasons. Arrows show fitted climatic predictors and axis labels the proportion of total variation represented. (b) Occurrence-frequency heatmap for the 33 taxa detected in at least three weeks. Cell colour gives the number of traps with detections in each week. The upper strip denotes meteorological season and the break separates the sampling periods.

The incidence-frequency heatmap showed the taxon-level patterns underlying temporal community variation (**Fig. 5b**). Recurrent taxa were generally detected more frequently during summer and autumn, with detections becoming markedly sparser in winter. The clearest period-specific pattern was observed for *Hanseniaspora valbyensis*, which was concentrated in winter 2022-2023, whereas *Magnusiomyces magnusii* was among the few taxa detected during winter in both sampling periods. Other taxa were less seasonal: *Debaryomyces prosopidis* occurred repeatedly throughout both sampling periods, while *Maudiozyma wufongensis* and *Lachancea fermentati* recurred mainly during warmer weeks. Thus, temporal compositional change reflected shifts in the occurrence of several taxa rather than replacement by a small number of strictly seasonal species.

### Thermal performance varies among yeast taxa and strains

Taxa differed in the collection-week mean temperatures associated with their incidence records, with several recovered predominantly below or above the long-term annual mean of 18.2°C (**Fig. 6a**). Four taxa spanning this distribution were selected for thermal phenotyping (**Fig. 6b**; **Supplementary Fig. 3**). *Saccharomyces cerevisiae* had the highest thermal optimum (T_opt_ ≈ 35.7°C) and warmest 80% thermal performance breadth (TPB_80_, 29.7–38.7°C), followed by *Lachancea fermentati* (T_opt_ ≈ 34.4°C; TPB_80_, 28.4–37.9°C). *Lachancea lanzarotensis* had a lower optimum (T_opt_ ≈ 24.3°C) and narrower TPB_80_ (19.5–27.2°C). Within *Magnusiomyces magnusii*, the October isolates F8H003 and F1H010 had higher estimated thermal optima (T_opt_ ≈ 29.7°C; TPB_80_, 24.1–32.9°C) than the four isolates recovered in December or January (T_opt_ ≈ 25.2°C; TPB_80_, 20.1–28.3°C). Among these selected isolates, thermal optima varied in the same direction as recovery temperatures.

**Fig. 6.**
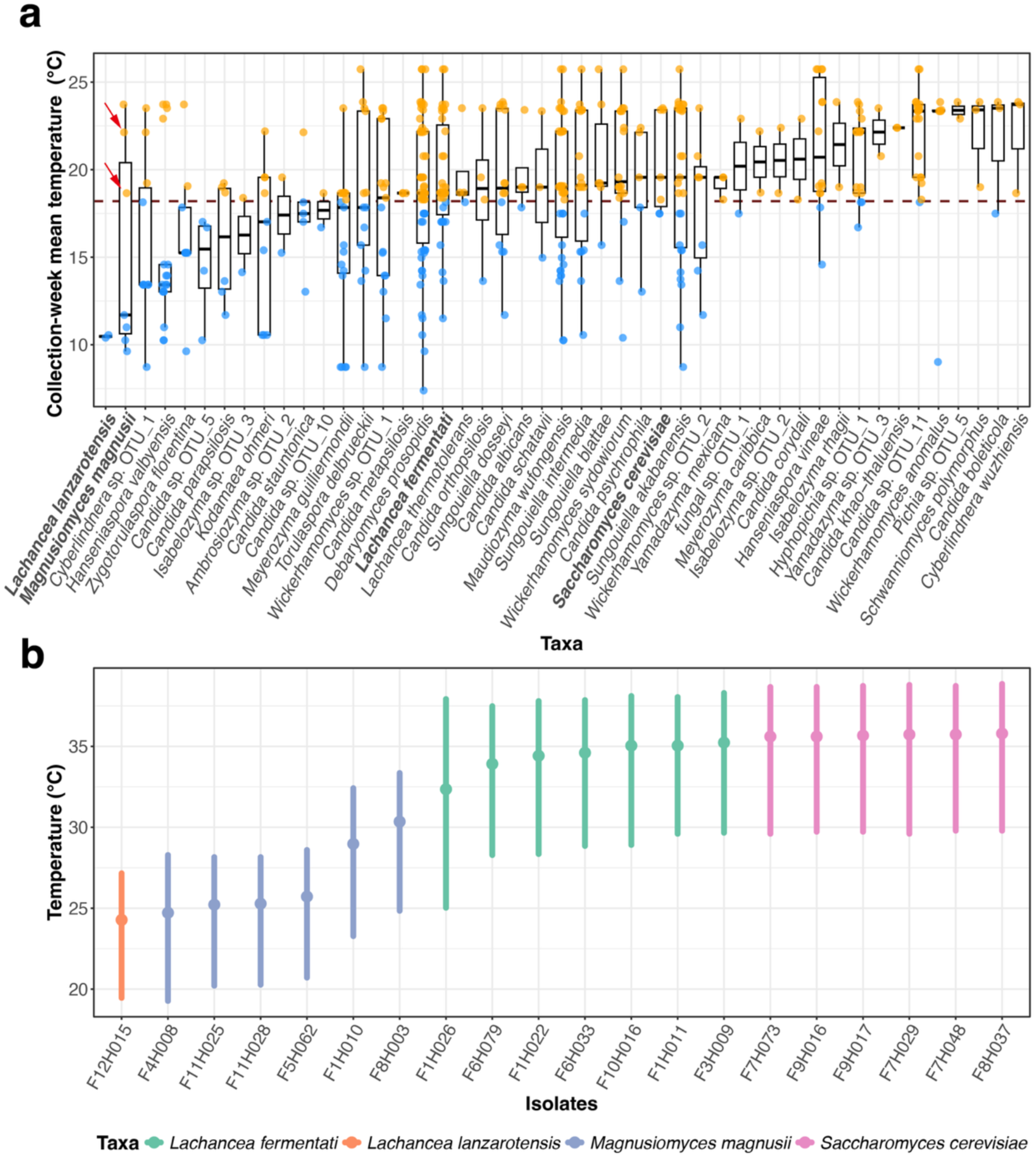
Collection-week temperatures and thermal performance of selected yeasts. (a) Weekly mean temperatures associated with incidence records for taxa detected at least twice. Blue and orange points denote temperatures below and above the long-term annual mean at Fushan (18.2°C), respectively. Box plots summarise each taxon’s distribution, bold labels mark phenotyped taxa and red arrows identify the two October *Magnusiomyces magnusii* isolates. (b) Estimated thermal optimum (T_opt_) and 80% thermal performance breadth (TPB_80_) for 20 isolates from four taxa. Points denote T_opt_, vertical lines denote TPB_80_ and colours identify taxa. Fitted thermal performance curves for individual isolates are shown in **Supplementary Fig. 3**.

## Discussion

Environmental surveys typically detect Saccharomycotina at low relative abundance^7,8^, whereas intensive weekly cultivation recovered a diverse assemblage containing many putatively novel ITS-designated lineages. Among-trap dissimilarity was dominated by turnover^14^, but neither measured topography nor geographic distance explained composition, and turnover was consistent with randomised expectations. By contrast, richness and composition varied seasonally with climate. Together with the correspondence between field recovery temperature and thermal performance among selected isolates, these patterns are consistent with climatic filtering of the culturable assemblage, although they do not establish causation or adaptation.

Forest inventories and global occurrence analyses indicate that Saccharomycotina diversity remains incompletely characterised^5,6,10–12,18,23^. Within 0.25 km², this survey recovered 56 described species and 77 putatively novel ITS lineages, while rarefaction of the primer-standardised dataset indicated further unsampled culturable diversity. Comparable cultivation surveys have reported similar or lower richness (**Supplementary Table S4**)^10–12,23^. East Asian surveys have also uncovered deeply divergent *Saccharomyces* populations and previously undescribed yeast species^9,23,24^. These findings underscore uneven sampling across Asia^25^. Recent taxogenomic analysis expanded *Torulaspora* from 10 to 22 recognised species, illustrating both the hidden diversity within familiar genera and the evidence needed to establish species boundaries^26^. By resolving community-level Saccharomycotina diversity within a single forest plot, this study complements population-focused studies of *Saccharomyces*^9,23,24^ and the global model of David et al., which modelled 186 described species at approximately 1-km resolution using heterogeneous occurrence records^5^. Direct richness comparisons therefore require comparable spatial grain, substrate, taxonomic coverage and detection methods^5,27^.

Temperature was the clearest environmental correlate of seasonal richness and composition, while thermal performance among selected isolates covaried with field recovery temperature. Temperature-dependent distributions of wild yeasts have been documented previously^19,22^, although seasonal patterns differ among habitats. Temperate forest soils can show a winter abundance maximum dominated by the psychrotolerant yeast *Tausonia pullulans*^15^, whereas an intertidal survey found highest diversity in autumn and lowest diversity in winter^28^. The Fushan pattern is consistent with temperature-related filtering, but incubation temperature also influences which taxa are recovered^27^. Field filtering and cultivation bias therefore cannot be separated here. Moreover, the retained climatic predictors explained only 6.4% of temporal compositional variation after adjustment, indicating that unmeasured variation in litter moisture, chemistry, resource availability and decomposition stage may also contribute to temporal turnover^29^.

Within *Magnusiomyces magnusii*, the contrast between two October and four winter isolates was consistent with strain-level thermal variation associated with recovery season, but limited strain and temporal replication preclude inference of seasonal selection or persistent warm- and cold-adapted lineages. A survey of 4,101 environmental yeast isolates similarly found extensive phenotypic variation among species but limited evidence that habitat consistently predicted within-species differentiation beyond particular taxa^30^. Resolving whether this pattern reflects stable population differentiation will require isolates from repeated warm and cold periods, common-environment phenotyping and genome-scale analyses.

Rarity dominated the primer-standardised ecological assemblage, with most taxa represented by a single incidence record or restricted to one trap. This pattern is consistent with the rare-biosphere framework^7,31^, and the prevalence of putatively novel lineages shows how incomplete occurrence data can affect diversity estimates^5^. Rare detection should not be equated with dispersal limitation and may reflect narrow niches, episodic activity, immigration, low abundance or low detection probability^14,31^. Neither distance decay nor greater-than-expected turnover supported dispersal limitation at this scale. Incomplete detection nevertheless prevents its exclusion. Recovery was also method-dependent. Isolation method and incubation temperature shape the taxa and phenotypes recovered^27,32,33^, while leaf-storage conditions can exert additional season-dependent effects^34^. The later use of ITS3 for isolates that failed full-length amplification illustrates primer-dependent recovery, and ITS does not resolve all fungal lineages uniformly^35^. The 98.5% threshold therefore defines operational units rather than species. Formal delimitation may require multilocus, genomic and phenotypic evidence^25^. We therefore treat the unidentified OTUs as putatively novel ITS lineages whose taxonomic status remains to be tested.

This study establishes subtropical forest leaf litter as a rich, underexplored reservoir of culturable Saccharomycotina yeasts and links community occurrence with climatic variation and isolate-level physiology. Its spatially and temporally explicit framework provides a basis for distinguishing transient recovery patterns from persistent ecological and evolutionary differentiation. Combining culture-independent detection, litter-level microclimate monitoring and genome-resolved phenotyping will help resolve how environmental, methodological and population-level processes generate forest yeast diversity.

## Materials and Methods

### Study site and trap deployment

The study was conducted in the Fushan Forest Dynamics Plot (FDP), located at the border of New Taipei City and Yilan County in northern Taiwan (24°45’N, 121°35’E; **Fig. 1a**). Fushan FDP is situated within a submontane evergreen broadleaf forest of the *Machilus*–*Castanopsis* vegetation zone. The plot spans 600 to 733 metres above sea level and is characterised by complex topography and mature evergreen broadleaf forest. Long-term climate records indicate a mean annual temperature of 18.2°C, mean annual precipitation of 4,271 mm and mean relative humidity of 95.1%^36^.

Fushan FDP serves as a long-term ecological research platform for investigating the processes shaping forest diversity and spatiotemporal dynamics in Taiwan. Since 2002, the site has supported phenological monitoring of plant flowering and fruiting in relation to climate change, together with quinquennial tree censuses of all woody stems within the plot^36^. As part of this infrastructure, 87 permanent seed traps (each 0.5 m²) were established to collect fallen flowers and seeds. Building on this existing system, we installed 20 circular leaf litter traps adjacent to 20 of these permanent seed traps (**Fig. 1b and 1c**). Each litter trap had an effective collection area of approximately 0.1 m² and was positioned to sample litter inputs from the surrounding canopy. The traps spanned hills, ridges, flat areas and streamside locations to capture local environmental heterogeneity.

### Sample collection and incubation

Leaf litter was collected weekly from the 20 litter traps (**Fig. 1b** and **1c**) during two sampling periods: 4 October 2022–8 February 2023 and 8 August 2023–15 February 2024. Each trap–week combination constituted one sample, yielding 20 samples per week (**Fig. 1d**). Samples were stored at 4°C immediately after collection and processed within two weeks. Trap–week combinations were retained as sampling units in the incidence matrix. Spatial analyses subsequently pooled detections by trap, whereas temporal analyses pooled detections by sampling week.

The amount of leaf litter collected per trap varied weekly, influenced by environmental factors such as wind and precipitation, and ranged from a few leaves to 150 pieces per sample. As the forest canopy at Fushan FDP is dominated by members of the Fagaceae, leaf litter from this family frequently constituted the majority of collected materials, with some samples containing up to 96% Fagaceae leaves (**Fig. 1e**). Fagaceae leaves were preferentially selected for incubation when present. For each sample, 8–12 leaf pieces were selected and divided between two 50 mL tubes, one for each incubation temperature. In the absence of Fagaceae leaves, leaf litter from other plant families was selected at random. Excess litter not used for incubation was discarded.

To maximise recovery of yeasts with different thermal preferences, each sample was incubated under two temperature regimes. After leaf material was transferred into Falcon tubes, 25 to 35 mL of high-sugar enrichment medium (HS; 10% glucose and 5% ethanol) was added to each tube^9^. Samples were incubated at either 20°C or 12°C, resulting in two parallel incubations per sample. Incubations at 20°C were conducted for 2 to 3 weeks, whereas samples incubated at 12°C were maintained for at least three weeks to recover slower-growing or cold-tolerant taxa.

### Yeast isolation and sequencing

Following incubation, samples were examined periodically and considered ready for isolation once a visible pellet had formed at the bottom of the Falcon tube. Leaf litter was removed without disturbing the pellet. Approximately 100 µL of the pellet suspension was then transferred onto yeast extract–peptone– dextrose (YPD) agar plates and streaked using a sterilised toothpick. Plates were incubated at room temperature for 2 to 5 days, after which 687 yeast-like colonies were selected based on colony morphology and re-streaked onto fresh plates to obtain pure cultures. The resulting isolates were maintained for downstream molecular identification.

Yeast isolates were identified by colony PCR targeting the internal transcribed spacer (ITS) region, followed by Sanger sequencing. Genomic DNA was extracted directly from single colonies using QuickExtract^TM^ according to the manufacturer’s instructions. PCR amplification was initially performed using the primer pair ITS1Fngs (GGT CAT TTA GAG GAA GTA A) and ITS4 (TCC TCC GCT TAT TGA TAT GC). From September 2023 onward, ITS3 (GCA TCG ATG AAG AAC GCA GC) was paired with ITS4 in cases where full-length ITS amplification was unsuccessful, to improve amplification success across diverse taxa, particularly those yielding weak or partial ITS amplicons. Briefly, a single colony was suspended in 10 µL of extraction buffer, vortexed and incubated at 65°C for 20 min, followed by 98°C for 5 min, then cooled to room temperature. Colony PCR was carried out in a 25 µL reaction containing 12.5 µL of PCR master mix, 1 µL of each primer, 9.5 µL of nuclease-free water, and 1 µL of extracted DNA. The PCR programme consisted of an initial denaturation at 95°C for 3 min, followed by 30 cycles of 95°C for 30 s, 52°C for 30 s, and 72°C for 1 min, with a final extension at 72°C for 5 min. PCR products were verified on 1% agarose gel, and successfully amplified samples were subjected to Sanger sequencing.

### Species classification, OTU clustering and sequence analysis

All ITS sequences obtained from yeast isolates were queried against the NCBI core_nt (version 20241216) database using BLASTn^37^. Species-level assignments required ≥98.5% sequence identity and ≥90% query coverage. Isolates that failed either threshold or matched reference sequences only at genus level were retained for OTU clustering without formal species-level designation.

Operational taxonomic units (OTUs) were defined using USEARCH v11.0^38^ by clustering sequences at 98.5% identity with a minimum alignment coverage of 90%. The centroid sequence represented each OTU. BLAST-based taxonomic assignments and OTU clustering results were integrated to establish the final classification. Clusters without species-level assignments but with clear affinity to known genera were designated as putatively novel OTUs and annotated at genus level. Ambiguous assignments were manually curated using BLAST scores, alignment coverage and phylogenetic context. Genome data or well-resolved reference sequences were used to refine species-level assignments where available. Taxonomic assignments and OTU classifications are summarised in **Supplementary Table S1**.

Phylogenetic analysis of novel *Hanseniaspora* OTUs was conducted using representative ITS sequences together with reference sequences of described species retrieved from NCBI GenBank. Sequences were aligned using MAFFT^39^ with the E-INS-i algorithm and trimmed with trimAl^40^, retaining alignment columns containing non-gap nucleotides in at least 30% of the sequences. A maximum-likelihood tree was reconstructed using IQ-TREE 3^41^ with the best-fit nucleotide substitution model selected by ModelFinder^42^. Branch support was assessed using 2,000 ultrafast bootstrap replicates and 1,000 SH-aLRT replicates, with BNNI correction applied to the ultrafast bootstrap analysis.

### Thermal phenotyping of selected yeast taxa

Four Saccharomycotina species were selected for targeted thermal phenotyping to span contrasting field recovery temperature patterns. Strains were selected across multiple traps and sampling conditions. To span within-species thermal variation, isolates from weeks differing by > 5°C in mean temperature were preferentially retained when multiple strains were available from the same trap. Thermal growth assays followed established protocols^24^ and were conducted at 10, 15, 20, 25, 30, 35, and 40°C. An additional assay at 27.5°C was performed for *Lachancea lanzarotensis*, which grew at 25°C but not at 30°C. Each isolate was tested in five biological replicates, with positive and negative controls included in triplicate, and all wells were randomly assigned within 96-well plates to minimise positional bias.

Yeast colonies were inoculated into 100 µL of YPD medium, vortexed, and incubated overnight at 25°C. Cultures were serially diluted to standardise cell density (wild strains: 10,000-fold; laboratory strains: 1,000-fold), and 15 µL of diluted culture was transferred into 135 µL of fresh YPD for optical density measurements. Growth was monitored at OD 595 nm using a TECAN Infinite M200 PRO plate reader. Measurements were recorded every 10 min for two days at moderate to high temperatures, and at hourly intervals for up to seven days at lower temperatures to accommodate slower growth.

Maximum growth rates were estimated using the gcplyr package^43^ and normalised for inter-plate variation using BY4741 as an internal reference. For each temperature, a plate-specific scaling factor was calculated as the ratio of the across-plate median BY4741 growth rate to the corresponding plate-specific median and applied to all isolates measured on that plate. Thermal performance curves were fitted using a custom implementation of the cardinal temperature model with inflection (CTMI) to estimate temperature-dependent growth parameters^44^. For each fitted curve, T_opt_ was the estimated temperature of maximum growth and TPB_80_ the temperature interval predicted to support at least 80% of that maximum.

### Statistical analyses

Analyses were conducted in R, with data processing and visualisation using readxl (https://CRAN.R-project.org/package=readxl), tidyverse^45^, reshape2^46^ and ggplot2^47^. Ecological analyses were restricted to isolates identified with ITS1Fngs/ITS4 to avoid bias introduced by the later use of ITS3/ITS4; ITS3/ITS4 identifications were retained only in the complete isolate inventory. Repeated recovery of the same taxon within a trap–week sample was scored once, producing a binary trap–week-by-taxon incidence matrix (**Supplementary Table S5**). Spatial analyses summed these detections across weeks for each trap (**Supplementary Table S6**), whereas temporal analyses summed detections across traps for each week (**Supplementary Table S7**). Matrix values therefore represent incidence or occurrence frequency, not environmental abundance.

Sampling completeness for the primer-standardised ecological dataset was assessed by rarefaction and extrapolation using iNEXT^48^. For whole-dataset rarefaction, the 497 taxon-incidence records were analysed as count data across the 124 taxa. For among-trap analyses, weekly detections were treated as the 47 incidence sampling units for each trap. Observed richness, incidence-based Chao2 and Shannon diversity calculated from weekly detection frequencies were obtained for each trap. The 20 × 124 trap-by-taxon matrix was converted to presence–absence for analyses of community composition.

Among-trap community composition was analysed using Jaccard dissimilarity. Standardised altitude, slope and distance to the nearest stream (**Supplementary Table S8**) were fitted as constraints in vegan::dbrda (https://CRAN.R-project.org/package=vegan); the overall model and marginal terms were tested using 9,999 unrestricted permutations. WGS84 trap coordinates were projected to a local metric coordinate system, and Euclidean distances were calculated between all trap pairs. One-sided Mantel tests with 9,999 permutations tested the a priori prediction that Jaccard dissimilarity increased with geographic distance, both across all taxa and among taxa detected at three or more traps. Jaccard dissimilarity was partitioned into turnover and nestedness-resultant components using betapart; mean values across the 190 non-independent trap pairs were treated descriptively. A curveball randomisation was applied separately to the complete 20 × 124 trap-by-taxon matrix and the recurrent 20 × 30 matrix, preserving the number of taxa per trap and the occurrence frequency of each taxon within each analysed matrix. Four independent chains generated 10,000 null matrices after burn-in. Two-sided empirical P values were calculated as min[1, 2 × min(Pgreater, Pless)] with +1 correction by comparing observed total dissimilarity, turnover and nestedness with their respective null distributions. The proportional contribution of turnover was also summarised for taxa detected in at least two, three and five traps.

All climatic variables, including temperature, precipitation, relative humidity, wind speed, and global solar radiation, were obtained from the Climate Observation Data Inquire Service of the Central Weather Administration of Taiwan and matched to sampling dates using records from automatic weather station C0AH90 (https://codis.cwa.gov.tw/StationData). Sampling weeks were assigned to seasons according to meteorological seasons in the Northern Hemisphere (**Supplementary Table S9**).

For temporal analyses, weekly Chao2 and Shannon indices were calculated from trap-level occurrence frequencies. Each climatic variable was tested in a separate linear mixed-effects model, with weekly Chao2 richness as the response, the climatic variable as a fixed effect and year–month as a random intercept. For community composition, the week-by-taxon occurrence-frequency matrix was converted to presence–absence and analysed by dbRDA of binary Jaccard dissimilarities among sampling weeks. Candidate environmental variables were grouped by ecological relevance and screened using single-variable dbRDA models. Within each group, the variable with the strongest explanatory performance was retained as the representative predictor. When only one usable variable was available, including cases in which alternatives contained excessive missing data, that variable was retained, whereas groups with no significant candidate variables were excluded from the final model (**Supplementary Table S3**). Overall and marginal terms were assessed by permutation tests.

## Data availability

Representative ITS sequences have been deposited in GenBank under accession numbers PZ490393–PZ490535, with corresponding strains listed in **Supplementary Table S1**. The binary trap–week-by-taxon incidence matrix, derived trap- and week-level occurrence-frequency matrices, trap metadata and matched weekly climatic records used in the ecological analyses are provided in **Supplementary Tables S5–S9**. All strains retained in the culture collection are available from the corresponding author upon reasonable request.

## Acknowledgements

IJT is funded by National Science and Technology Council, Taiwan (Grant NSTC 113-2628-B-001-002 and 114-2628-B-001-014) and Academia Sinica (Grant AS-IA-113-L04).

## Contributions

IJT conceived the study. CHCY provided the Fushan Forest Dynamics Plot as the research platform. IJT, YCL, CHCY and CHT established the litter-trap sampling system, and CHT conducted weekly field sampling. WTC performed experiments with assistance from CH, PWS, and CJY. YCY performed OTU clustering. WTC conducted diversity and spatiotemporal analyses with guidance from IJT, CTT, PJK, and CHCY. WTC and IJT wrote the manuscript with comments from all authors.

