## Supplementary Information for "High culturable diversity and climate-associated seasonal dynamics of Saccharomycotina yeasts in subtropical forest leaf litter"

**Supplementary Document**

**Supplementary Tables**

**Supplementary Table S1.** Taxonomic assignments and OTU classifications of yeast isolates recovered in this study.

This table summarizes the taxonomic assignments and OTU classifications of 143 yeast OTUs recovered in this study. Taxonomic assignments and OTU clustering were performed following the procedures described in the Materials and Methods. For taxa with highly similar reference sequences, cases involving multiple plausible BLAST hits or shared cluster centroids were manually curated, and relevant decisions were recorded in the “Additional note” column. The representative strain was assigned as the centroid strain when available; otherwise, the strain with the highest sequence identity was selected, with random selection applied when identity values were identical.

**Supplementary Table S2.** Summary of sampling, yeast recovery, identification, and final datasets.

**Supplementary Table S3.** Summary of environmental variables used in the ecological analyses.

The table lists all predictor and variables included in the different statistical models and their corresponding P-values.

**Supplementary Table S4.** Comparison of Saccharomycotina diversity and novelty across forest ecosystems.

The table summarizes reported estimates of yeast diversity, novelty, and sampling scale from this study and selected forest ecosystems in different studies.

**Supplementary Table S5.** Incidence matrix of culturable Saccharomycotina across all litter-trap samples.

Rows represent individual trap-by-sampling-time combinations (Trap × Time), columns represent taxa, and cell values indicate species incidence (0/1). This matrix was used as the input dataset for community ecological analyses.

**Supplementary Table S6.** Trap-level incidence matrix of culturable Saccharomycotina.

Rows represent individual litter traps, with occurrences pooled across all sampling times. Columns represent taxa, and cell values indicate the number of sampling occasions in which each species was detected in each trap. This matrix was used for spatial ecological analyses.

**Supplementary Table S7.** Time-level incidence matrix of culturable Saccharomycotina.

Rows represent individual sampling occasions, with occurrences pooled across all litter traps. Columns represent taxa, and cell values indicate the number of litter traps in which each species was detected during each sampling occasion. This matrix was used for temporal ecological analyses.

**Supplementary Table S8.** Geographic and environmental characteristics of the 20 litter-trap sampling locations in the Fushan Forest Dynamics Plot.

**Supplementary Table S9.** Weekly climatic variables matched to the 47 valid sampling weeks in the Fushan Forest Dynamics Plot.

**Supplementary Figures**

**Supplementary Fig. 1** ITS-based phylogeny of *Hanseniaspora* showing the positions of novel OTUs recovered in this study. Novel OTUs were highlighted in red. Colored circles at the branch nodes indicate bootstrap support categories, with green representing high support (≥ 90), orange representing moderate support (70–89), and red representing low support (< 70).

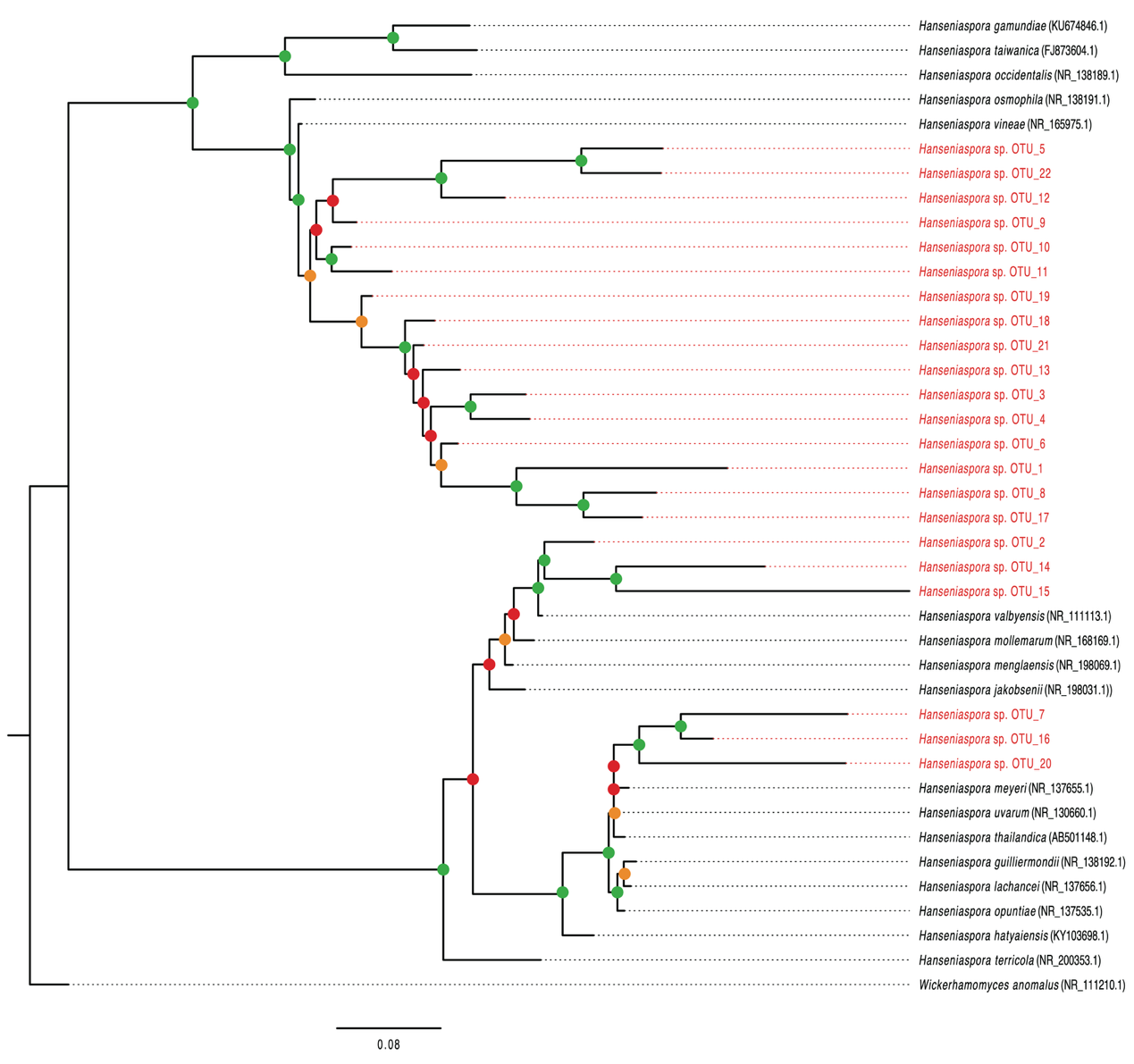

**Supplementary Fig. 2** Linear mixed models showing the relationships between Chao2-estimated richness and other environmental variables, including weekly mean temperature, maximum temperature, minimum temperature, cumulative precipitation, relative humidity, minimum relative humidity, wind speed, maximum gust, and global radiation. Each point represents one sampling week.

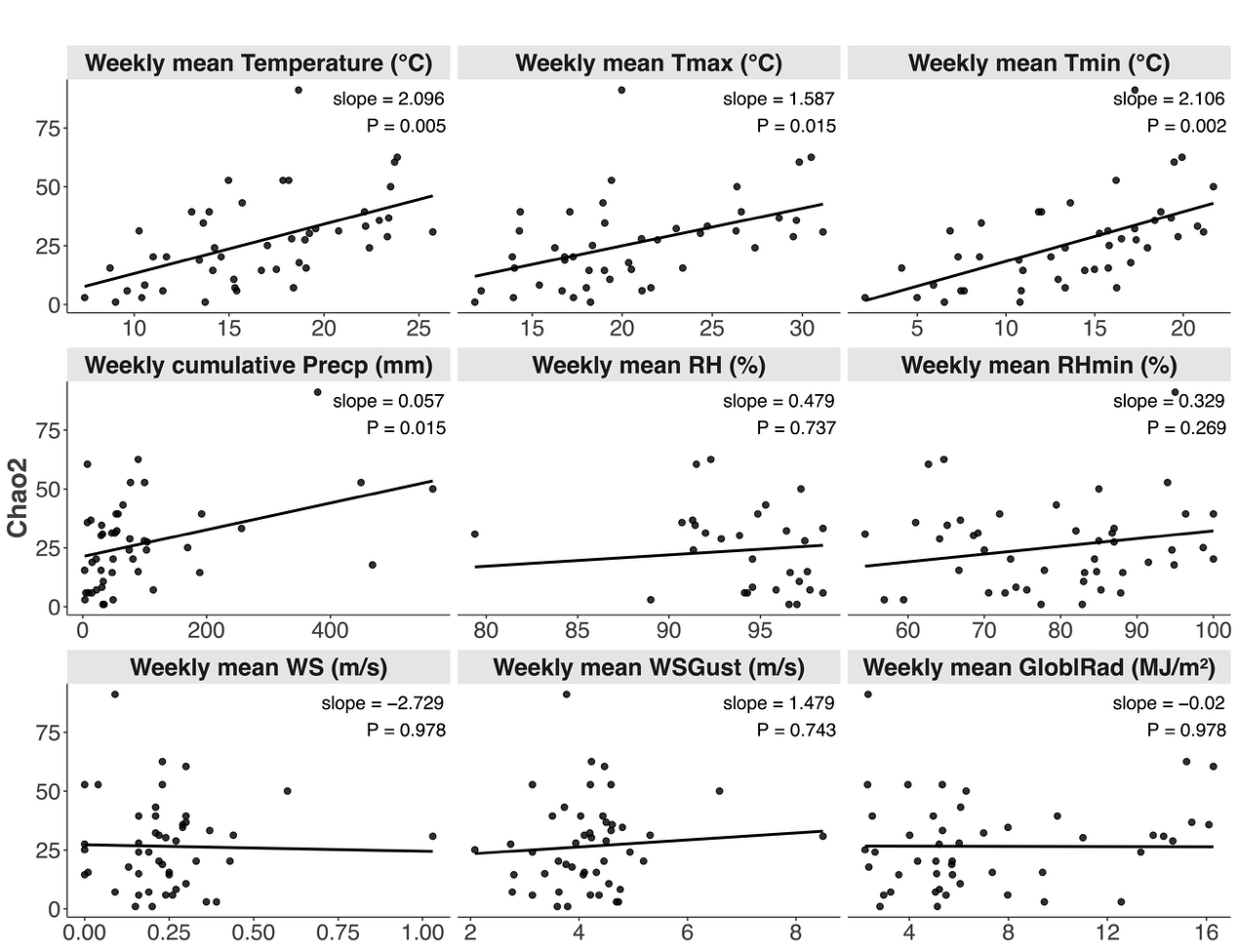

**Supplementary Fig. 3** Thermal performance curves of representative yeast isolates. Maximum growth rate as a function of temperature for representative isolates of *Saccharomyces cerevisiae*, *Lachancea fermentati*, *Magnusiomyces magnusii*, and *Lachancea lanzarotensis*. Each curve represents the fitted Cardinal Temperature Model with Inflection (CTMI) for a single isolate.

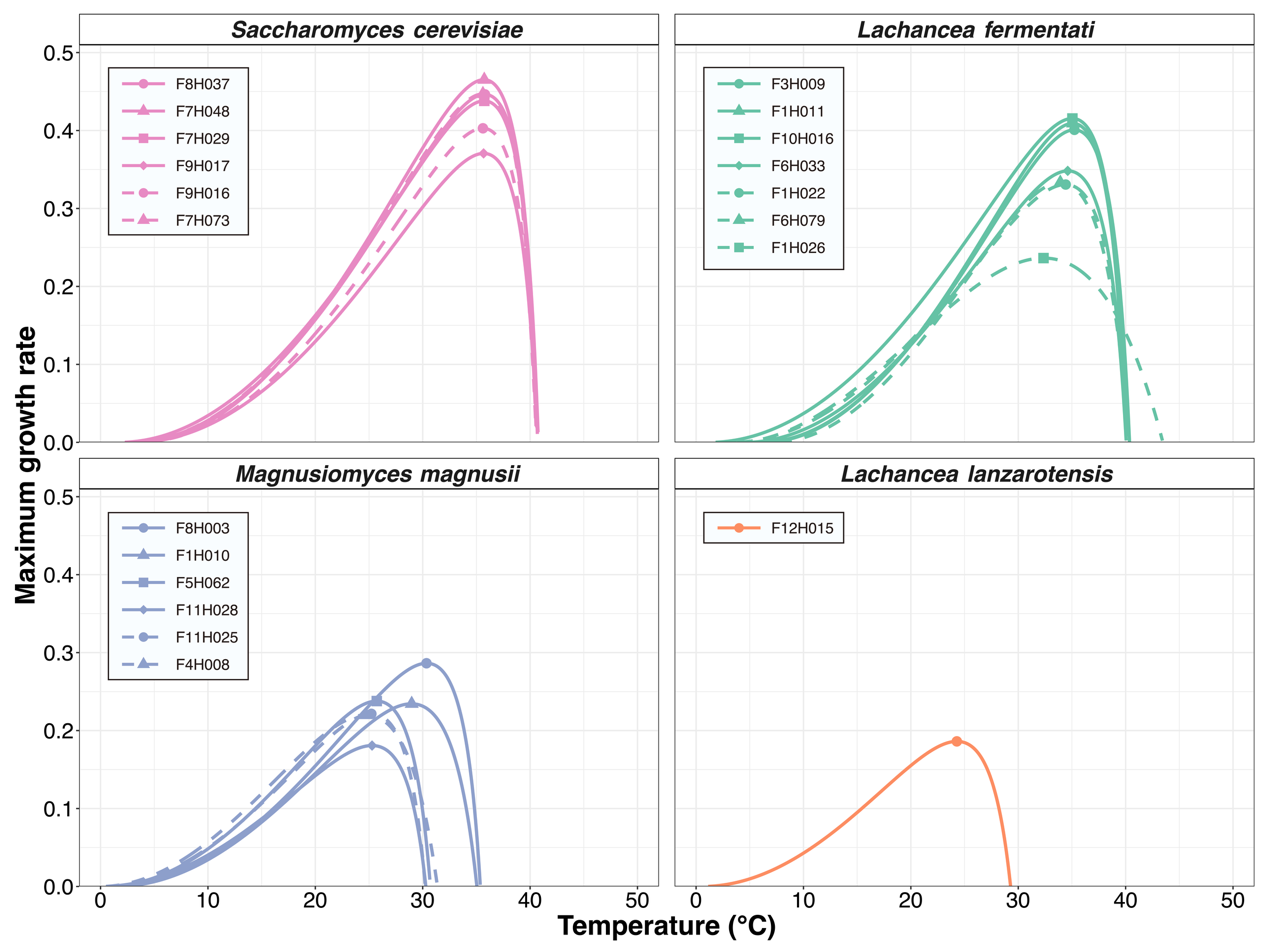
